# Investigating the physical effects in bacterial therapies for avascular tumors

**DOI:** 10.1101/683839

**Authors:** Pietro Mascheroni, Michael Meyer-Hermann, Haralampos Hatzikirou

## Abstract

Tumor-targeting bacteria elicit anticancer effects by infiltrating hypoxic regions, releasing toxic agents and inducing immune responses. Although current research has largely focused on the influence of chemical and immunological aspects on the mechanisms of bacterial therapy, the impact of physical effects is still elusive. Here, we propose a mathematical model for the anti-tumor activity of bacteria in avascular tumors that takes into account the relevant chemo-mechanical effects. We consider a time-dependent administration of bacteria and analyze the impact of bacterial chemotaxis and killing rate. We show that active bacterial migration towards tumor hypoxic regions provides optimal infiltration and that high killing rates combined with high chemotactic values provide the smallest tumor volumes at the end of the treatment. We highlight the emergence of steady states in which a small population of bacteria is able to constrain tumor growth. Finally, we show that bacteria treatment works best in the case of tumors with high cellular proliferation and low oxygen consumption.

## 1. Introduction

Cancers display huge variability between different patients and even in the same patient. Nonetheless, cancer cells share a finite set of hallmarks such as sustained proliferation, invasion and metabolic reprogramming, which shape their behavior in solid tumors (Hanahan and Weinberg, 2011). Among other hallmarks, tumor cells are known to recruit new blood vessels to sustain their proliferation, in a process known as *tumor angiogenesis* (Folkman, 1971). This neovasculature is generally altered in terms of architecture and morphology of the vessels, leading to poor perfusion of certain areas of the tumor (Carmeliet and Jain, 2000). Hypoxic regions are thus created and maintained during tumor development, concurring to the progression of cancer cells towards malignant phenotypes (Vaupel and Mayer, 2007). Moreover, low nutrient levels can lead to cell quiescence, a situation in which tumor cells delay metabolic activities and become less sensitive to standard chemotherapies (Challapalli et al., 2017). Such hypo-perfused areas are generally associated with poor patient outcome but, on the other hand, could be exploited for tumor targeting (Wilson and Hay, 2011). The same hypoxic areas provide indeed a niche for bacteria to colonize the tumor and exert a therapeutic action (Forbes, 2010; Zhou et al., 2018). The use of bacteria for cancer therapy dates back hundreds of years, with doctors reporting tumor regression in several patients (Kramer et al., 2018). However, such treatments also caused some fatalities and the limited understanding of the therapeutic mechanisms of action shifted research efforts towards other strategies - especially radiotherapy (Kramer et al., 2018). In the last few years the use of live bacteria for cancer treatment has regained interest, and several bacterial strains have been tested in animal models and even advanced to clinical trials (Torres et al., 2018). Nevertheless, clinical development of such therapies is still facing significant issues due to infection-associated toxicities and incomplete knowledge of infection dynamics (Kramer et al., 2018; Zhou et al., 2018). As much research was focused on the immune responses after bacteria administration, a clear picture of the interaction between cancer and bacterial cells is still lacking.

Mathematical modeling emerges as a promising candidate to assist the understanding of bacterial therapy mechanism of action in cancer. Mathematical models have been applied in the context of cancer to elucidate its progression and treatment (Byrne, 2010; Altrock et al., 2015). Recent examples combining experimental and modeling work in bacterial therapies are given in (Kasinskas and Forbes, 2006; Jean et al., 2014; Hatzikirou et al., 2017; Suh et al., 2018), featuring *in vitro* as well *in vivo* experiments.

Here we describe a mathematical model for bacteria-based cancer therapy within avascular tumors, focusing on the influence of physical effects on therapy outcomes. Such effects are present in every biological system but are often concealed by the complexity of the interactions between molecular and cellular players. Here we show through a simple mathematical model that these effects take an important part in bacterial therapies and are able to influence their outcomes. The model is formulated in the context of mixture theory, a framework with a long history of applications to biological problems - see for example Ambrosi and Preziosi (2002); Breward et al. (2001, 2002, 2003); Byrne and Preziosi (2003); Chaplain et al. (2006); Preziosi and Tosin (2009) and the recent reviews of Siddique et al. (2017); Pesavento et al. (2017). Our aim is to evaluate the impact of bacterial chemotaxis and anti-tumor activity on cancer cells, using spheroids as a prototype of avascular tumors. We consider bacterial administration after full formation of the spheroid, when hypoxic areas are present. We describe the effects of the treatment on the behavior of the spheroid constituents, e.g. tumor cells and bacteria volume fractions, at different time points and over the spheroid radius.

The remainder of the paper is organized as follows. In Section 2 we describe the mathematical model and its derivation. In Section 3 we present model results, analyzing the impact of different model parameters. Finally, in Section 4 we discuss the biological implications of the results and suggest new research directions.

## 2. Materials and Methods

We propose a mathematical model describing the impact of bacterial cells on tumor spheroid growth. The model is based on mixture theory, a continuum theory that allows to describe the chemo-mechanical interactions between different tissue components. We follow the approach discussed in Preziosi (2003); Byrne (2012) and, specifically, adapt the derivation in Boemo and Byrne (2019) to our problem. In the following we present the final form of the equations, leaving the full derivation in the Supplementary Information.

We describe the tumor as being composed of three main constituents (or *phases* in the language of mixture theory): tumor cells (TCs), bacteria and extracellular material. The variables referring to these quantities will be identified by the indexes c, b and f, respectively. We also consider the presence of a nutrient, i.e. oxygen, diffusing over the spheroid domain. The model equations are derived by applying conservation of mass and linear momentum to each phase, and enforcing the saturation constraint (i.e. all the space in the spheroid is occupied by the phases, there are no voids). Then, we close the model by imposing suitable constitutive assumptions regarding the material properties of the phases and their interaction terms.

### 2.1. Model equations

In the following we will be interested in the case of tumor spheroids, for which the assumption of spherical symmetry applies. The problem reduces to the set of Partial Differential Equations (PDEs):

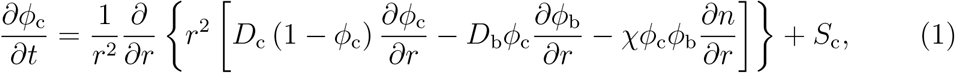

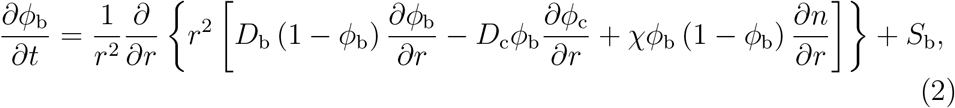

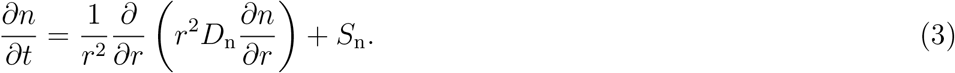

Here, *ϕ*_c_, *ϕ*_c_ and *n* are the tumor cell and bacteria volume fractions and normalized nutrient concentration, respectively. These quantities depend on the radial coordinate *r* ∈ [0, *R*] and time *t* ∈ [0, *t_f_*]. In addition, *D*_i_ are the phases motility coefficients (i=c,b), *D*_n_ the nutrient diffusion coefficient, and *χ* the bacterial chemotactic coefficient. The mass exchange terms *S*_i_ (i=c,b,n), regulating the transfer of mass between the different components, will be detailed in the next subsection. Note that we do not solve explicitly for *ϕ*_f_ (i.e. the volume fraction of extracellular material) since this quantity can be obtained as *ϕ*_f_ = 1 − *ϕ*_c_ − *ϕ*_b_ due to the saturation constraint (see the Supplementary Information). We model growth of the spheroid as a free-boundary problem, in which the outer tumor radius *r* = *R*(*t*) moves with the same velocity as the TC phase,

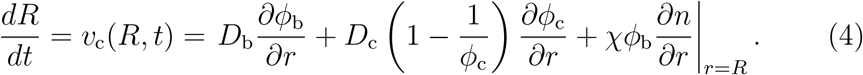

Finally, we define a set of boundary and initial conditions to close the differential problem in equations (1)–(3). Due to the problem symmetry no-flow boundary conditions are enforced at the spheroid center, whereas we fix the values of TC volume fraction, bacterial volume fraction and normalized nutrient concentration on the spheroid boundary:

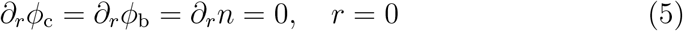

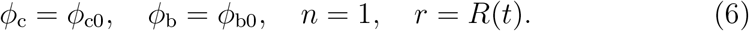

We assume a uniform initial tumor volume fraction *ϕ*_c0_ = 0.8 across the spheroid (Byrne and Preziosi, 2003) and, to model bacteria administration, we consider a time dependent value for the bacterial volume fraction at the spheroid outer radius:

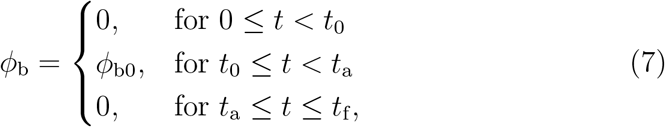

where *ϕ*_b0_ is the administered volume fraction of bacteria, *t*_0_ is the time of administration and *t*_a_ its duration. Regarding the initial conditions, we consider a spheroid devoid of bacteria and displaying a uniform TC volume fraction and nutrient concentration over its radius:

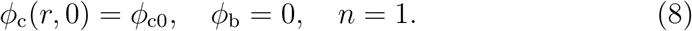

Finally, we prescribe an initial spheroid radius, i.e. *R*(0) = 90 *μ*m. The equations of the model are discretized through the Finite Element Method and solved using the commercial software COMSOL Multiphysics (COMSOLAB).

### 2.2. Choice of mass exchange terms

To formulate the mass exchange terms in equations (1)–(3) we assume the following assumptions (see Figure 1):

**A1** TCs proliferate when oxygen is available. As soon as the latter decreases below a critical threshold, they stop proliferating and start necrosis (Chaplain et al., 2006; Gerlee and Anderson, 2007; Agosti et al., 2018).
**A2** Bacteria compete with TCs for space and exert an anti-tumor effect by a variety of mechanisms (e.g. by realising toxins and therapeutic agents, or stimulating an immune response). (Forbes, 2010; Osswald et al., 2015; Torres et al., 2018; Zhou et al., 2018).
**A3** Bacteria die when oxygen is above a critical threshold and thrive in hypoxic conditions (*anaerobic* bacteria) (Toley and Forbes, 2011; Phaiboun et al., 2015; Osswald et al., 2015).
**A4** TCs consume oxygen provided by the culture medium (Matzavinos et al., 2009; Grimes et al., 2014).

**Figure 1:**
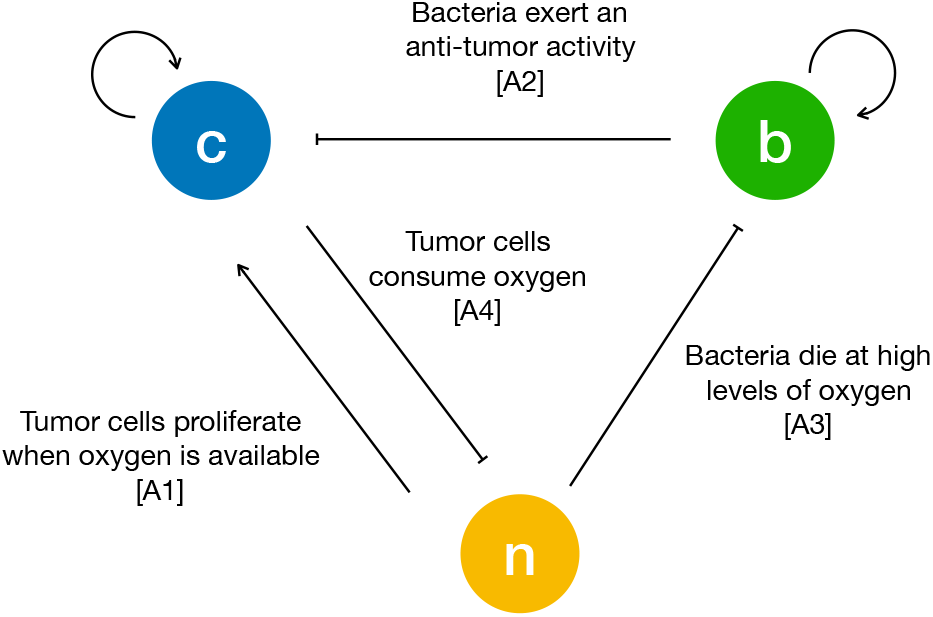
Schematic of the interactions between tumor cells (*c*), bacteria (*b*) and oxygen (*n*). The arrows are drawn according to the biological hypotheses detailed in the main text.

The resulting mass exchange terms read:

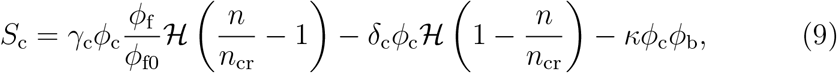

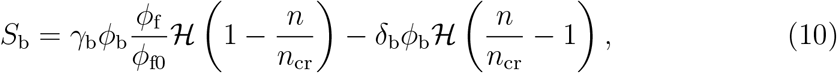

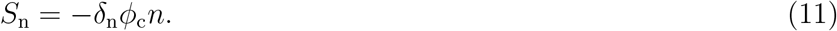

Here *γ*_i_ and *δ*_i_ are the proliferation and death rate of the i-th phase respectively (i = c, b), whereas *δ*_n_ is the oxygen consumption rate. In addition, *ϕ*_*f*0_ is the initial volume fraction of extracellular material and we indicate with 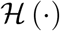 a smooth version of the step function, and with *n*_cr_ the critical oxygen value below which hypoxic conditions develop. Finally, we do not consider a specific form for the anti-tumor effect of bacteria and introduce an effective TC killing rate *κ* in the equation for *S*_c_.

### 2.3. Model parametrization

The parameters used in the model simulations are reported in Table 1. In the following we will compare model results with a set of published experiments on the U87 glioma cell line. We take the TC proliferation rate from the available data provided from the Bioresource Core Facility of the Physical Sciences-Oncology Center (PBCF, 2012), whereas we select the TC death rate in accordance to the estimate in (Kolokotroni et al., 2011; Martínez-González et al., 2012). The work in (Toley and Forbes, 2011) provides a value for the bacterial motility coefficient and proliferation rate in *in vitro* cellular aggregates. Regarding bacterial proliferation, (Gibson et al., 2018) supply a similar value through an analysis of bacterial doubling times. We estimate the bacterial death rate from (Phaiboun et al., 2015), in which cellular death dynamics are quantified under starvation at different bacteria densities. Finally, we use the values in (Schaller and Meyer-Hermann, 2005; Matzavinos et al., 2009; Grimes et al., 2014; Colombo et al., 2015; Alfonso et al., 2016) for the oxygen diffusion coefficient and consumption rate in tumor tissues. When carrying out the simulations, we vary the chemotactic coefficient in the interval [0, 8.64 × 10^−1^]mm^2^d^−1^. Since it was not possible to find in the literature an estimate for the chemotactic coefficient of bacteria in tissues, we considered the value of *χ* in bacterial solutions (Ford et al., 1991; Lewus and Ford, 2001) and divided it for the ratio between the motility coefficient in solution and in tissue - about 100, (Ford et al., 1991; Lewus and Ford, 2001). Since we do not consider a specific mechanism for the anti-tumor activity of bacteria, we select the killing rate *κ* to be in the interval [0, 5] d^−1^, i.e. spanning characteristic times between several days and a few hours. Finally, we fit the parameter for the critical oxygen concentration from the above mentioned experiments. The value that we found is similar to the one reported in (Gerlee and Anderson, 2007; Agosti et al., 2018).

**Table 1:** Summary of the parameter values considered in the model simulations.

| Parameter | Value | Description | Reference |
| --- | --- | --- | --- |
| $D_c$ | $0.5 \text{ mm}^2 \text{d}^{-1}$ | TC motility coefficient | (Colombo et al., 2015) |
| $\gamma_c$ | $0.48 \text{ d}^{-1}$ | TC proliferation rate | (PBCF, 2012) |
| $\delta_c$ | $0.5 \text{ d}^{-1}$ | TC death rate | (Martínez-González et al., 2012) |
| $D_b$ | $0.05 \text{ mm}^2 \text{d}^{-1}$ | Bacterial motility coefficient | (Toley and Forbes, 2011) |
| $\gamma_b$ | $15 \text{ d}^{-1}$ | Bacterial proliferation rate | (Gibson et al., 2018) |
| $\delta_b$ | $0.24 \text{ d}^{-1}$ | Bacterial death rate | (Phaiboun et al., 2015) |
| $D_n$ | $100 \text{ mm}^2 \text{d}^{-1}$ | Oxygen diffusion coefficient | (Matzavinos et al., 2009) |
| $\delta_n$ | $8640 \text{ d}^{-1}$ | Oxygen consumption rate | (Colombo et al., 2015) |
| $\chi$ | $[0, 0.864] \text{ mm}^2 \text{d}^{-1}$ | Bacterial chemotactic coefficient | estimated |
| $\kappa$ | $[0, 5] \text{ d}^{-1}$ | Bacterial killing rate | model specific |
| $n_{cr}$ | 0.58 | Critical oxygen concentration | calibrated |

## 3. Results

### 3.1. Model calibration on spheroid experiments

We start the analysis by considering the growth of a spheroid suspended in culture medium, in the absence of bacteria. We compare the results of the model with the data for radial growth of U87 tumor spheroids available from Mascheroni et al. (2016). We use the model to fit the critical oxygen concentration parameter *n*_cr_, keeping all the other quantities as defined in Table 1. Figure 2 shows a good agreement between the model and the experiments, over all the growth curve. The model is able to reproduce the two phases of spheroid growth usually described in the literature (Conger and Ziskin, 1983; Sutherland, 1988; Vinci et al., 2012). The spheroid radius (see Figure 2**A**) displays a first stage of rapid increase, followed by a saturation phase. This behavior is detailed in Figures 2**B,C**, showing the evolution of the tumor volume fraction and oxygen concentration over the spheroid radius at different time points. The tumor volume fraction, i.e. *ϕ*_c_, increases over the spheroid at early time points (Figure 2**B**). Then, as TCs consume oxygen to proliferate, its concentration decreases in the centre of the aggregate (Figure 2**C**). When the oxygen level drops below the critical threshold *n*_cr_ (dashed line in Figure 2**C**), TCs stop proliferating and die. This results in a decrease of *ϕ*_c_ in the spheroid core, displayed at longer times in Figure 2**B**. Close to saturation, the amount of cells that proliferate is balanced by the number of cells that die, turning into extracellular material. Therefore, even if cell growth continues to take place in the outer rim of the spheroid, it is not enough to advance the spheroid front, which reaches a steady state. These results match qualitatively what is observed in the experimental (Landry et al., 1982; Montel et al., 2011; Grimes et al., 2014; Sarkar et al., 2018)and modeling (Ward and King, 1999; Byrne and Preziosi, 2003; Ambrosi and Mollica, 2004; Schaller and Meyer-Hermann, 2005; Mascheroni et al., 2016; Boemo and Byrne, 2019) literature for tumor spheroids and will serve as a basis for the discussion in the next sections.

**Figure 2:**
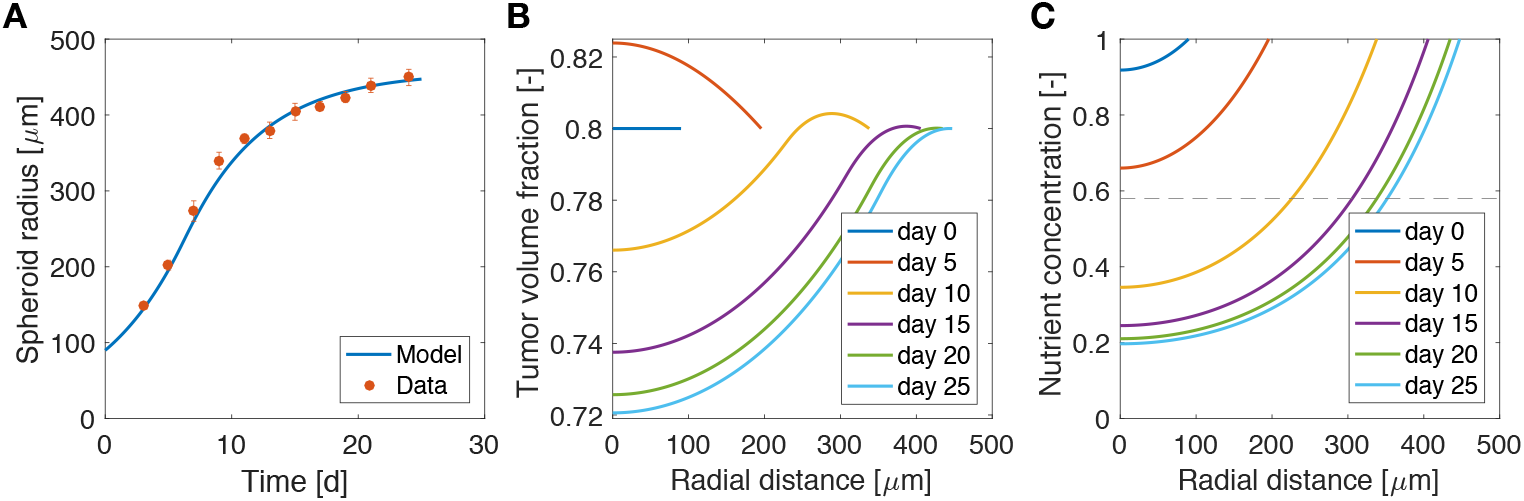
Calibration of the model on tumor spheroid data. **A** Comparison between model results and experimental data for the spheroid growth curve. The experimental points are taken from (Mascheroni et al., 2016) and represent the growth of U87 spheroids. Dots are mean values and bars standard deviation of the measurements. Tumor volume fraction (**B**) and oxygen concentration (**C**) at different times of spheroid growth. The dashed line in the last plot displays the critical oxygen concentration.

### 3.2. Administration of bacteria leads to tumor remission but not eradication

Figure 3 shows the influence of bacterial therapy on tumor spheroid composition for an example case. We evaluate the effects of adding bacteria to the culture medium after the spheroid is fully formed, i.e. when hypoxic regions have developed. In particular, we select an administration time of *t*_0_ = 26d and a treatment duration of *t*_a_ = 2d. We consider an intermediate value for both the bacterial chemotactic coefficient and killing rate (*χ* = 0.432mm^2^d^−1^, *κ* = 2.5d^−1^). As shown by the low TC volume fraction in Figure 3**A** at later times, bacteria administration leads to spheroids less populated by TCs. This space is occupied by bacteria (Figure 3**B**), which thrive in the hypoxic region located in the spheroid core. After bacterial therapy the spheroid shrinks and is less populated by cancer cells. This leads to higher values of oxygen concentration at the center of the aggregate, as displayed in Figure 3**C**. Finally, Figure 3**D** shows the evolution of TC (*V*_c_) and bacteria (*V*_b_) volumes over time. These quantities are calculated as

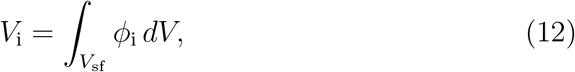

where the integral is performed over the spheroid volume *V*_sf_ (i=c,b). At early time points, *V*_c_ is in a phase of fast growth, since the nutrient is available throughout the spheroid and no bacteria are present. After administration, there is a fast increase of bacteria volume together with a rapid decrease of TC volume. At later time points the system evolves toward a steady state in which both bacteria and TCs coexist in the tumor aggregate. Even though the TCs are not completely removed, the spheroid persists in an equilibrium state, where an asymptotic size is kept for long times.

**Figure 3:**
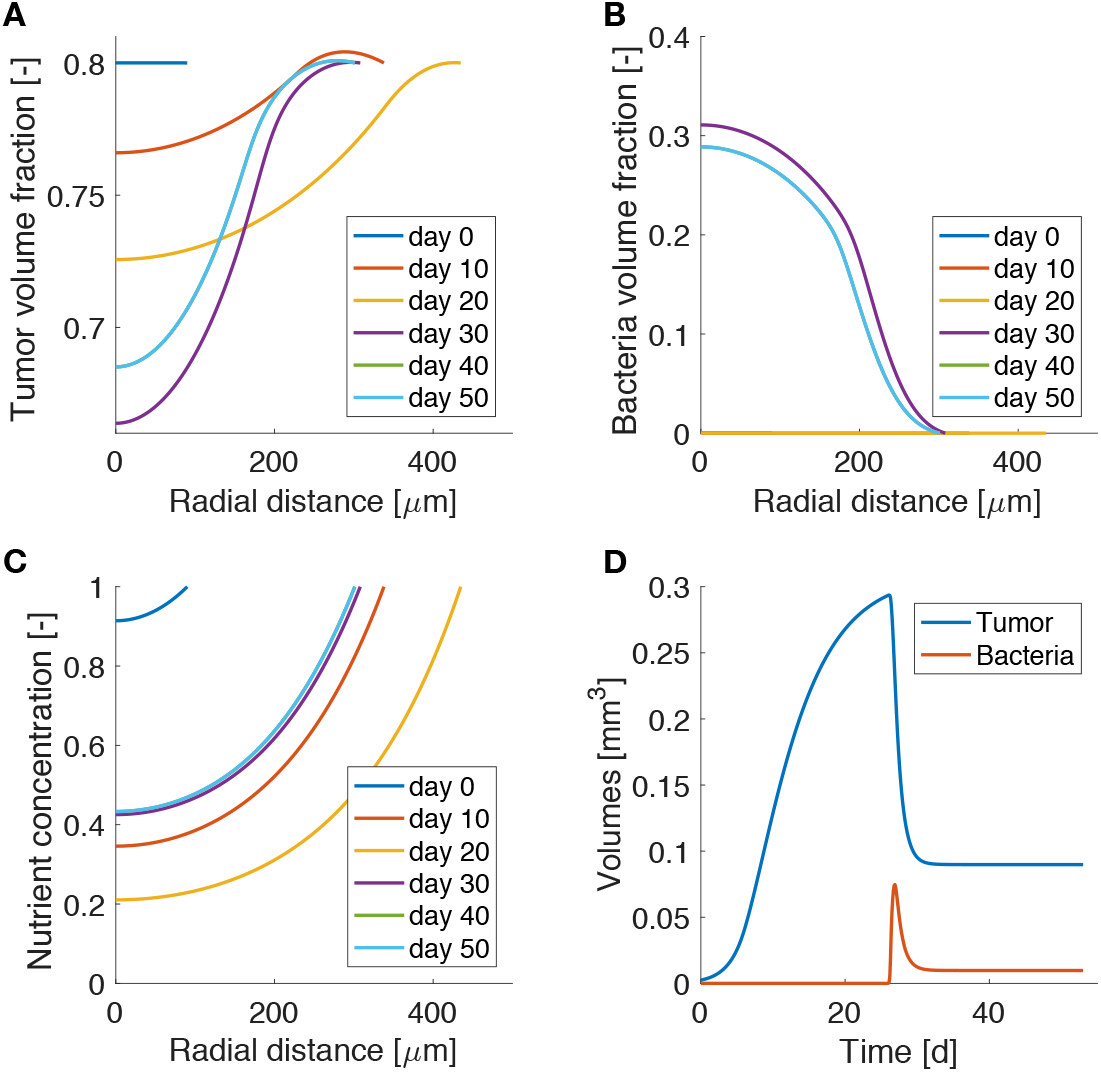
Model results for bacteria administration to tumor spheroids. Spatio-temporal evolution of tumor (**A**) and bacteria (**B**) volume fractions and oxygen concentration (**C**). **D** Temporal evolution of tumor and bacteria volumes in the spheroid.

### 3.3. High chemotaxis allows for maximal reduction of tumor size

We investigated the impact of different bacterial chemotactic and anti-tumor strengths on spheroid composition at the end of the simulations, i.e. at day 50 (Figure 4). We found that the highest reduction in tumor volume is obtained for the highest values of the chemotactic coefficient and killing rate, as shown in Figure 4**A**. On the other hand, highly chemotactic bacteria without an anti-tumor activity lead to the highest tumor volume. Interestingly, the tumor is never completely eradicated over all the explored parameter sequence. A similar result is obtained for the bacteria volume at the end of the simulations (Figure 4). Here, no matter the strength of chemotaxis or anti-tumor activity, bacterial cells are always present in the final spheroid volume. High bacterial volumes are present for high chemotactic coefficients, whereas high killing rates lead to small bacterial volumes independent of the chemotactic strength. Indeed, even though the tumor volume considerably varies over the chemotactic space for high killing rates, the bacterial volume is almost independent of this quantity (see Figure S1 in the Supplementary). Finally, Figures 4**C** and 4**D** show the temporal variation of tumor and bacterial volumes for two extreme cases occurring for high chemotaxis and low (Case 1) or high (Case 2) killing rate. The first plot shows that after the administration of bacteria the tumor volume is reduced, even in the absence of anti-tumor activity. The two populations in the spheroid reach an equilibrium at later times, with bacteria representing a significant portion of the spheroid. In the second case, the high anti-tumor activity of the bacteria is responsible for a sharp decrease of the tumor population, leading also to oscillations in the TC volume. Although bacteria now constitute a small part of the overall spheroid volume, they are still able to keep the tumor size under control.

**Figure 4:**
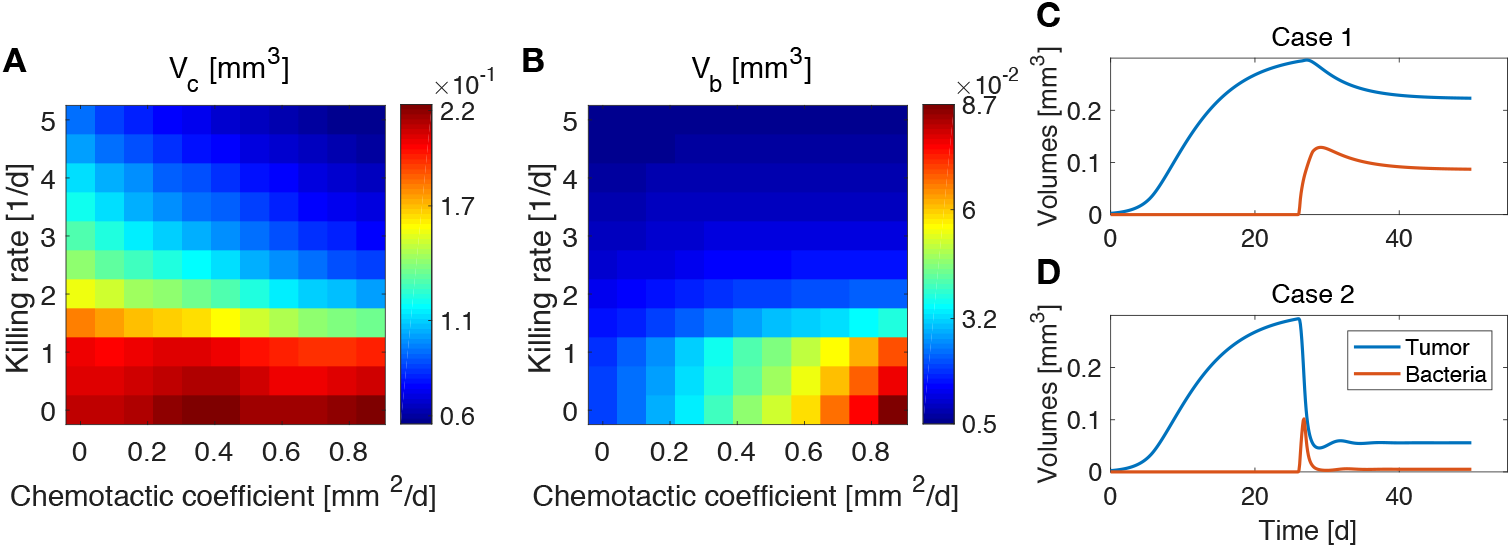
Influence of bacteria chemotactic coefficient and anti-tumor activity on tumor (**A**) and bacteria (**B**) volumes at the end of the simulations (day 50). Temporal evolution of tumor and bacteria volumes for a high chemotactic coefficient and a low (Case 1, **C**) and high (Case 2, **D**) killing rate.

### 3.4. Highly proliferating and low oxygen consuming tumors are mostly benefited from bacterial therapy

The results obtained in the previous subsection are insensitive of the administration time *t*_0_, the duration of the administration *t*_a_ and the administered bacteria volume fraction *ϕ*_b0_, even for large variations of these parameters (see Supplementary Figures S2-S4). This made us investigate whether the steady states reached at the end of the simulations and displayed in Figure 4 were therefore a function of the mechanisms regulating the tumor/bacteria dynamics. To check this hypothesis we simulated the behavior of TCs with a lower or higher proliferation and oxygen consumption rates with respect of the one shown in Figure 4. We report our findings in Figures 5 and 6. We considered a variation of ±50% with respect to the nominal value of the parameters in Table 1, and labeled the cases using the plus or minus in the superscript accordingly. All the other parameters keep the nominal values. We evaluated the spheroid response in terms of relative tumor reduction by introducing the quantity:

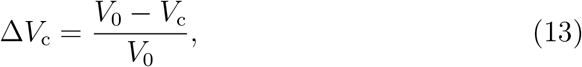

where *V*_c_ is the final tumor volume and *V*_0_ the tumor volume at the time of bacteria administration. We also analyzed the relative bacteria volume at the end of the simulation, plotting the ratio of bacteria volume *V*_b_ to the total spheroid volume *V*_t_.

**Figure 5:**
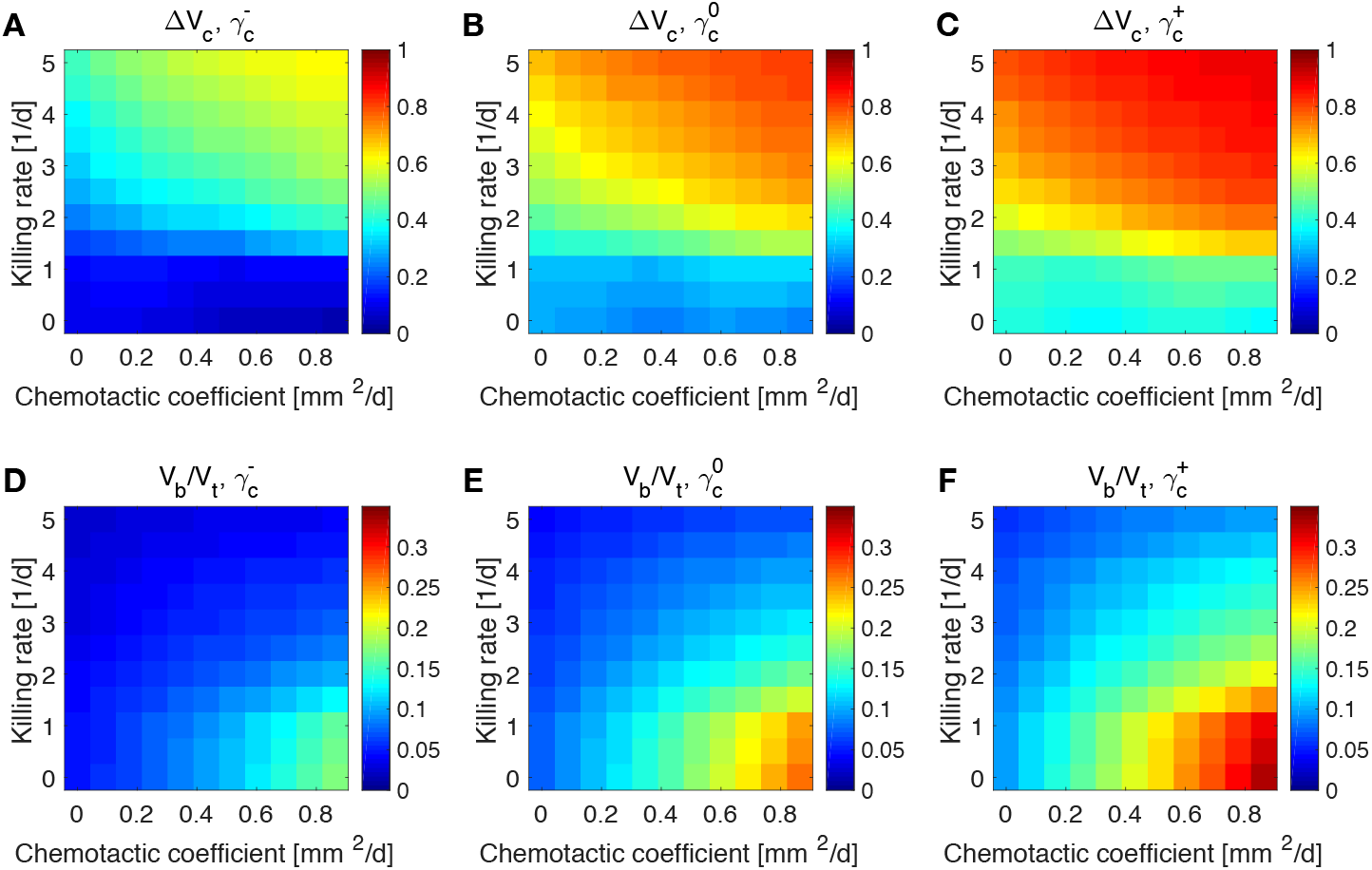
Impact of tumor proliferation rate on tumor and bacteria volumes at the end of the simulations (day 50). Relative tumor volume change and relative bacteria volume for low (**A, D**), nominal (**B, E**) and high (**C, F**) tumor cell proliferation rate.

**Figure 6:**
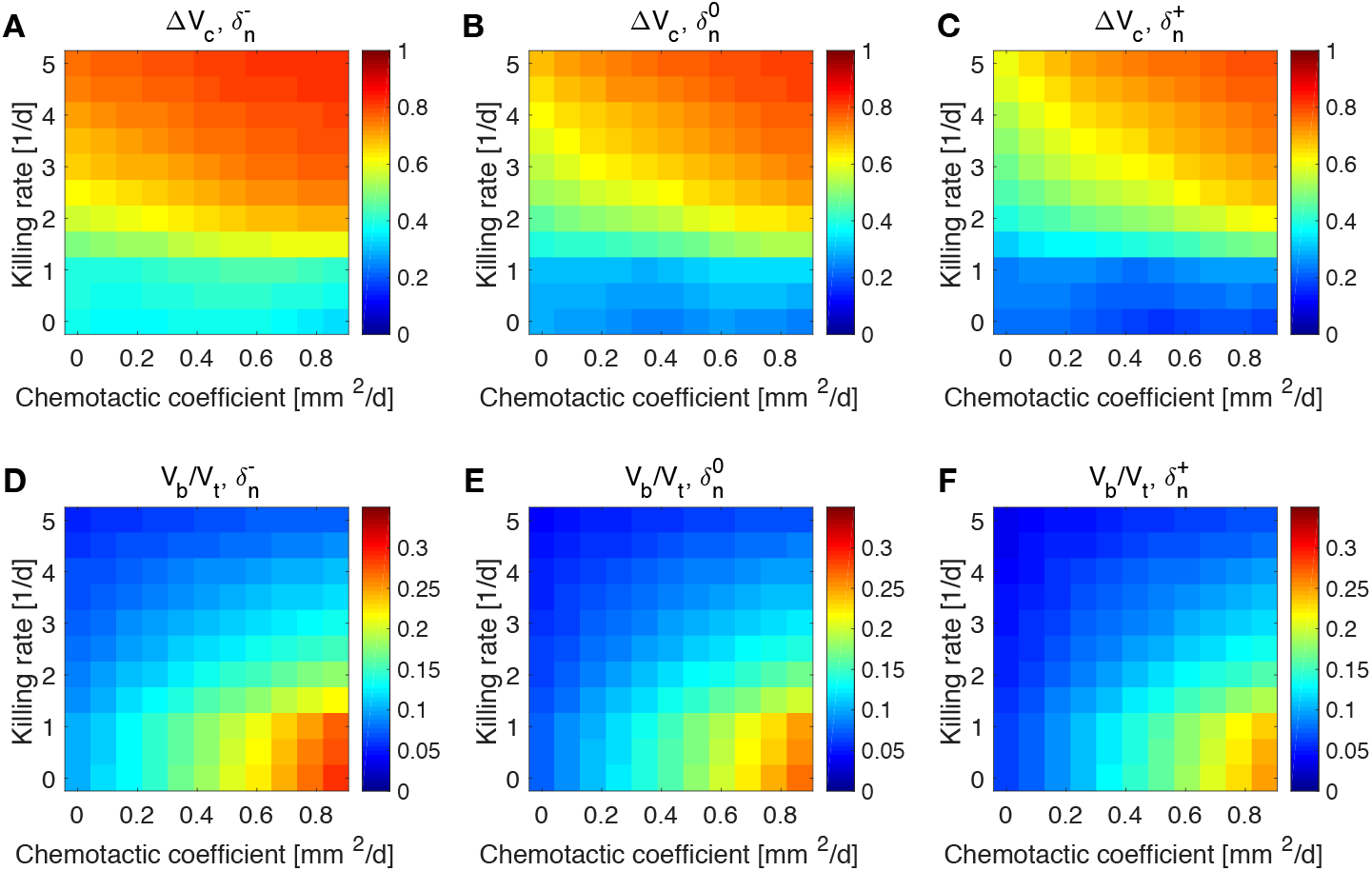
Impact of tumor oxygen consumption rate on tumor and bacteria volumes at the end of the simulations (day 50). Relative tumor volume change and relative bacteria volume for low (**A, D**), nominal (**B, E**) and high (**C, F**) tumor cell proliferation rate.

Tumors in which cells proliferate at a higher rate display the highest tumor reductions (Figure 5**A-C**). This is particularly true for the treatment with bacteria characterized by high chemotaxis and killing rate. Highly proliferative tumors are the ones that also show higher colonization by bacteria, as displayed in Figures 5**D-F**. Treatments with high chemotactic bacteria with low killing rates provide the highest relative bacteria volumes. Low oxygen consumption by TCs leads to results similar to highly proliferative tumors (6). Again, treatment using bacteria with high chemotaxis and high killing rate produces the best results in terms of tumor reduction. Regarding the final bacterial content, both high and low oxygen consuming tumors show considerable bacteria colonization. As before, the relative bacteria volume is higher for highly chemotactic bacteria with low anti-tumor activity. Even though highly proliferative and low oxygen consuming TCs originate the highest final spheroid volumes (Figure S5), they benefit the most from bacteria treatment and display the higher final bacteria content.

## 4. Discussion

We proposed a mathematical model to study the influence of bacteria treatment on avascular tumor growth. We considered anaerobic bacteria which thrive in hypoxic environments and actively migrate towards nutrient deprived regions in solid tumors. The model was calibrated to reproduce published tumor spheroid data and then used to evaluate the impact of bacteria chemotaxis and killing rate on spheroid response.

Model results show preferential bacteria accumulation in the hypoxic spheroid core, with tumor cells more localized towards the external spheroid surface. In general, highly chemotactic bacteria possessing increased anti-tumor activity provide the highest tumor reduction after treatment. On the other hand, high chemotaxis but low anti-tumor activity lead to smaller tumor reduction but higher bacteria colonization at the end of the simulations. When varying the tumor parameters, we found that bacteria treatment works best for highly proliferative and low oxygen consuming tumors.

For simplicity, we considered a general effective anti-tumor activity of TCs by bacteria without focusing on specific mechanisms, e.g. cytotoxic agents, prodrug-converting enzymes, etc. (Torres et al., 2018; Zhou et al., 2018; Kramer et al., 2018). Such treatment modalities could be incorporated by extending the model, to provide a more accurate description of the therapeutic action. Moreover, we focused on tumor spheroids, an *in vitro* approximation of avascular tumors. As such, they lack all the interactions between the tumor and its immune environment. On the one hand, this approach allows to investigate the mutual dynamics of bacteria and tumor cells without external influences, but on the other including the cross-talk between bacteria and the components of the immune system would be a fundamental step to address questions coming from *in vivo* tumors. Following (Boemo and Byrne, 2019), we modeled the mechanical response of cells and bacteria in the simplest way considering the phases as inviscid fluids. Although this description is still able to qualitatively describe the experimental results, more detailed constitutive assumptions for the mechanical behavior of the phases would lead to new insights into the interactions between bacteria and TCs in the aggregate (Sciumè et al., 2013; Giverso et al., 2015; Ambrosi et al., 2017; Mascheroni et al., 2018; Fraldi and Carotenuto, 2018; Giverso and Preziosi, 2019). We also considered ideal spherical spheroids to reduce the mathematical problem to one dimension. Even if the qualitative results will be maintained in a three-dimensional geometry, adopting the latter will be crucial to translate the model to *in vivo* situations.

In this modeling approach, space competition between bacteria and tumor cells arises naturally from the conservation of mass and momentum imposed by the governing equations. As no void regions are allowed into the spheroid, when cells move or die one of the model components automatically fills the space. Bacteria and TCs compete for space in the spheroid and the expansion of the tumor becomes limited, especially when the anti-tumor activity of bacteria is strong. However, for increasing values of the chemotactic coefficient and low values of the killing rate, bacteria localize predominantly in the spheroid core and displace TCs to the outer region of the spheroid. Both types of cell can proliferate in each of the two spheroid areas (hypoxic for spheroids, well-oxygenated for TCs), giving rise to high spheroid volumes and considerable bacteria colonization.

As a matter of fact, chemotaxis could be a target for bacteria-based anticancer therapies and diagnostic tools. For example, TCs that become restricted to outer spheroid areas after administration of highly chemotactic bacteria are more oxygenated and could benefit from standard chemotherapeutic or radiation treatments in the context of synergistic treatments (Zhou et al., 2018). We highlight that this is an example showing that mathematical models could help to identify situations when TC sensitization to therapies might be possible - see also (Owen et al., 2004; Kim et al., 2013; Michor and Beal, 2015; Mascheroni et al., 2017). On the other hand, highly chemotactic bacteria could be used as tracers to identify necrotic regions in spheroid, exploiting their targeting efficiency. Moreover, the simulations show the existence of steady states in which a small population of bacteria is in dynamical equilibrium with cancer cells, leading to tumor size control over time. All these mechanisms arise as a pure physical effect from the competition for space between cancer and bacteria cells and could be optimized to obtain the highest tumor volume reduction or bacteria colonization. Currently, even though researchers are aware of the benefits coming from active bacteria migration towards hypoxic regions in tumors (Forbes, 2010; Kramer et al., 2018), this knowledge has not been efficiently exploited in the clinical trials carried out so far (Torres et al., 2018).

Finally, we point out three straightforward developments that emerge from the findings of this work. Firstly, our theoretical results advocate for experiments with tumor spheroids. With such a simplified experimental setup, several bacterial strains could be tested on different cancer cell lines to validate model findings. Secondly, one could think about extending the model to consider different bacterial administration schedules. The duration of bacteria administration, the time of administration and single vs. multiple dosing could be investigated to determine the optimal conditions for this kind of treatment. Lastly, the tight coupling between the dynamics of TCs and bacteria in terms of regulating their reciprocal environment could be addressed via mathematical models, in order to control the bacterial infection or identify the optimal timing of the therapy.

## Conflict of Interest Statement

The authors declare that the research was conducted in the absence of any commercial or financial relationships that could be construed as a potential conflict of interest.

## Author Contributions

PM, HH and MMH contributed conception and design of the study; PM derived the model and performed the computational analysis; PM wrote the first draft of the manuscript; HH and MMH wrote sections of the manuscript. All authors contributed to manuscript revision, read and approved the submitted version.

## Funding

MMH and HH gratefully acknowledge the funding support of the Helmholtz Association of German Research Centers - Initiative and Networking Fund for the project on Reduced Complexity Models (ZT-I-0010). HH and PM acknowledge the funding support of MicMode-I2T (01ZX1710B) by the Federal Ministry of Education and Research (BMBF).

## Acknowledgements

We thank Dr. Raimondo Penta (University of Glasgow) for the help with the COMSOL simulations.

