## Supplementary Information for "Investigating the physical effects in bacterial therapies for avascular tumors"

### 1. Model derivation

Model derivation is carried out in the framework of mixture theory, following the approach discussed in Preziosi (2003); Byrne (2012). Specifically, we adapt the derivation in Boemo and Byrne (2019), which deals with a mixture model for macrophage-based therapies in tumor spheroids, to our problem. We consider a mixture constituted by cancer cells (c), bacteria (b) and extracellular material (f) - *phases*, in the following. We also include the presence of a nutrient (n), i.e. oxygen, in our description. Following (Preziosi, 2003; Byrne, 2012), we first write the balance of mass for each phase:

$$\partial_t \phi_{\mathrm{i}} + \mathrm{div}\,(\phi_{\mathrm{i}} \boldsymbol{v}_{\mathrm{i}}) = S_{\mathrm{i}}, \qquad (1)$$

in which $\phi_{\mathrm{i}}$, $\boldsymbol{v}_{\mathrm{i}}$ and $S_{\mathrm{i}}$ are the volume fraction, velocity and mass exchange term related to the i-th phase (i = c, b, f). Note that Equation (1) implicitly assumes that the phases have the same constant mass density. In the following we will also assume that the mixture is closed with respect to mass, so that mass can only be converted from one phase to the other, i.e. $S_{\mathrm{f}} = -S_{\mathrm{c}} - S_{\mathrm{b}}$.

*Correspondence:
 (M.M.H.);
 (H.H.)

In mixture theory velocity fields are determined by considering the mechanical response of the phases to mutual interactions Ambrosi and Preziosi (2002). Neglecting inertial effects, as usually done for growth phenomena (Preziosi, 2003; Byrne, 2012), the balance of linear momentum can be written as:

$$\mathrm{div}\left(\phi_{\mathrm{i}}\boldsymbol{\sigma}_{\mathrm{i}}\right) + \sum_{\mathrm{i}\neq\mathrm{j}} \boldsymbol{m}_{\mathrm{ij}} + p\,\mathrm{grad}\left(\phi_{\mathrm{i}}\right) = \boldsymbol{m}_{\mathrm{i}}. \tag{2}$$

Here $\boldsymbol{\sigma}_{\mathrm{i}}$ is the stress tensor of the i-th phase, $\boldsymbol{m}_{\mathrm{ij}}$ represent the forces exerted on the i-th phase by the j-th phase, and $\boldsymbol{m}_{\mathrm{i}}$ describes an external force acting on the i-th phase (i, j = c, b, f). Note that, for the action-reaction principle, $\boldsymbol{m}_{\mathrm{ij}} = -\boldsymbol{m}_{\mathrm{ji}}$. Finally, the terms $p\,\mathrm{grad}\left(\phi_{\mathrm{i}}\right)$ represent interfacial effects between phases, with $p$ being the interfacial pressure (Byrne, 2012). In this modeling framework, $p$ emerges as a Lagrange multiplier due to the saturation constraint

$$\sum_{\mathrm{i}=\mathrm{c,b,f}} \phi_{\mathrm{i}} = 1, \tag{3}$$

meaning that we assume that there are no empty spaces within the mixture (Preziosi, 2003; Byrne, 2012).

We conclude the set of governing laws by stating an equation for the normalized nutrient concentration $n$ in the mixture, i.e. the tumor:

$$\partial_t n = D_{\mathrm{n}}\mathrm{div}\left(\mathrm{grad}\, n\right) + S_{\mathrm{n}}, \tag{4}$$

in which $D_{\mathrm{n}}$ is the nutrient diffusion coefficient and $S_{\mathrm{n}}$ represents the nutrient mass exchange with the model phases.

#### *1.1. Constitutive relationships*

We close the model by selecting suitable constitutive assumptions. First, we assume that the interaction terms $\boldsymbol{m}_{\mathrm{ij}}$ depend linearly on the relative phase velocities (Preziosi, 2003; Byrne, 2012):

$$\boldsymbol{m}_{\mathrm{ij}} = -\mu\phi_{\mathrm{i}}\phi_{\mathrm{j}}\left(\boldsymbol{v}_{\mathrm{i}} - \boldsymbol{v}_{\mathrm{j}}\right), \tag{5}$$

with the same linearity constant $\mu$ for all the phases (i = c, b, f). We consider only a single external force $\boldsymbol{m}_{\mathrm{b}}$ acting on bacteria. This term describes bacteria chemotaxis following spatial hypoxic gradients and models active cell migration towards waste products from dying cancer cells (Forbes, 2010; Toley and Forbes, 2011). We assume a linear relationship,

$$\boldsymbol{m}_{\mathrm{b}} = \phi_{\mathrm{b}} \chi_{\mathrm{b}} \mathrm{grad}\, n, \tag{6}$$

in which $\chi_{\mathrm{b}}$ describes the strength of chemoattraction.

Following Breward et al. (2001, 2002); Byrne (2012); Boemo and Byrne (2019) we consider the phases as inviscid fluids and associate an interfacial pressure to each of them. For simplicity, we take the pressure in the extra-cellular material to be equal to that in the fluid surrounding the spheroid, $p$. The stress tensors in Equation (2) are defined such that the interfacial pressure of each phase is given by the pressure in the extracellular material plus a correction term, specific to its phase (Boemo and Byrne, 2019):

$$\boldsymbol{\sigma}_{\mathrm{f}} = -p\boldsymbol{I}, \tag{7}$$

$$\boldsymbol{\sigma}_{\mathrm{b}} = -(p + \pi_{\mathrm{b}})\boldsymbol{I}, \tag{8}$$

$$\boldsymbol{\sigma}_{\mathrm{c}} = -(p + \pi_{\mathrm{c}})\boldsymbol{I}, \tag{9}$$

where $\boldsymbol{I}$ is the identity tensor. The ratio $\pi_{\mathrm{i}}/\mu$ characterizes the movement of the i-th phase in the mixture and is generally identified as the phase motility coefficient $D_{\mathrm{i}}$ (i = c, b) (Boemo and Byrne, 2019). In the following we will also define $\chi = \chi_{\mathrm{b}}/\mu$ as the bacterial chemotactic coefficient.

#### *1.2. Spherical symmetry, initial and boundary conditions*

After recasting Equation (1) in spherical symmetry and after imposing the saturation constraint in Equation (3), we have:

$$v_{\mathrm{c}}\phi_{\mathrm{c}} + v_{\mathrm{b}}\phi_{\mathrm{b}} + v_{\mathrm{f}}\phi_{\mathrm{f}} = 0, \tag{10}$$

in which $v_{\mathrm{i}}$ is the radial velocity of the i-th phase (i = c, b, f). Substituting Equations (5), (7)-(9) and (10) in Equation (2) we obtain for the radial velocities:

$$v_{\mathrm{c}} = D_{\mathrm{b}} \frac{\partial \phi_{\mathrm{b}}}{\partial r} + D_{\mathrm{c}} \left(1 - \frac{1}{\phi_{\mathrm{c}}}\right) \frac{\partial \phi_{\mathrm{c}}}{\partial r} + \chi \phi_{\mathrm{b}} \frac{\partial n}{\partial r}, \tag{11}$$

$$v_{\mathrm{b}} = D_{\mathrm{b}} \left(1 - \frac{1}{\phi_{\mathrm{b}}}\right) \frac{\partial \phi_{\mathrm{b}}}{\partial r} + D_{\mathrm{c}} \frac{\partial \phi_{\mathrm{c}}}{\partial r} - \chi \left(1 - \phi_{\mathrm{b}}\right) \frac{\partial n}{\partial r}, \tag{12}$$

after summing over the phases in Equation (2) to express $p$ as a function of the other model quantities. Substituting Equations (11)-(12) in (1) and rewriting the system in spherical symmetry leads to the final model equations:

$$\frac{\partial \phi_{\mathrm{c}}}{\partial t} = \frac{1}{r^2}\frac{\partial}{\partial r}\left\{r^2\left[D_{\mathrm{c}}\left(1-\phi_{\mathrm{c}}\right)\frac{\partial \phi_{\mathrm{c}}}{\partial r} - D_{\mathrm{b}}\phi_{\mathrm{c}}\frac{\partial \phi_{\mathrm{b}}}{\partial r} - \chi\phi_{\mathrm{c}}\phi_{\mathrm{b}}\frac{\partial n}{\partial r}\right]\right\} + S_{\mathrm{c}}, \tag{13}$$

$$\frac{\partial \phi_{\mathrm{b}}}{\partial t} = \frac{1}{r^2}\frac{\partial}{\partial r}\left\{r^2\left[D_{\mathrm{b}}\left(1-\phi_{\mathrm{b}}\right)\frac{\partial \phi_{\mathrm{b}}}{\partial r} - D_{\mathrm{c}}\phi_{\mathrm{b}}\frac{\partial \phi_{\mathrm{c}}}{\partial r} + \chi\phi_{\mathrm{b}}\left(1-\phi_{\mathrm{b}}\right)\frac{\partial n}{\partial r}\right]\right\} + S_{\mathrm{b}}, \tag{14}$$

$$\frac{\partial n}{\partial t} = \frac{1}{r^2}\frac{\partial}{\partial r}\left(r^2 D_{\mathrm{n}}\frac{\partial n}{\partial r}\right) + S_{\mathrm{n}}. \tag{15}$$

Note that we do not solve for $\phi_{\mathrm{f}}$ since it can be obtained as $\phi_{\mathrm{f}} = 1-\phi_{\mathrm{c}}-\phi_{\mathrm{b}}$ through Equation (3).

### 2. Supplementary Figures

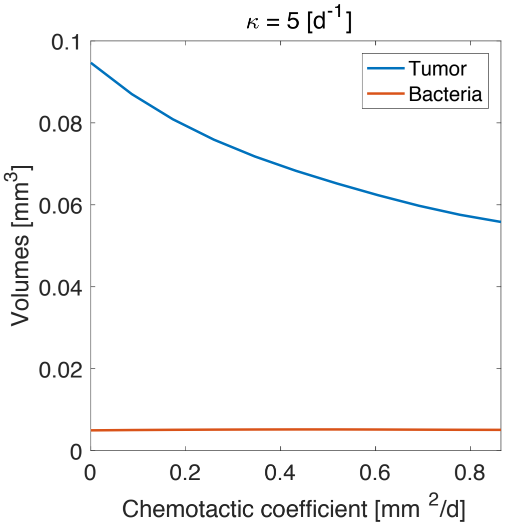


Figure S1: Influence of bacteria chemotactic coefficient on tumor and bacteria volume for high anti-tumor activity.

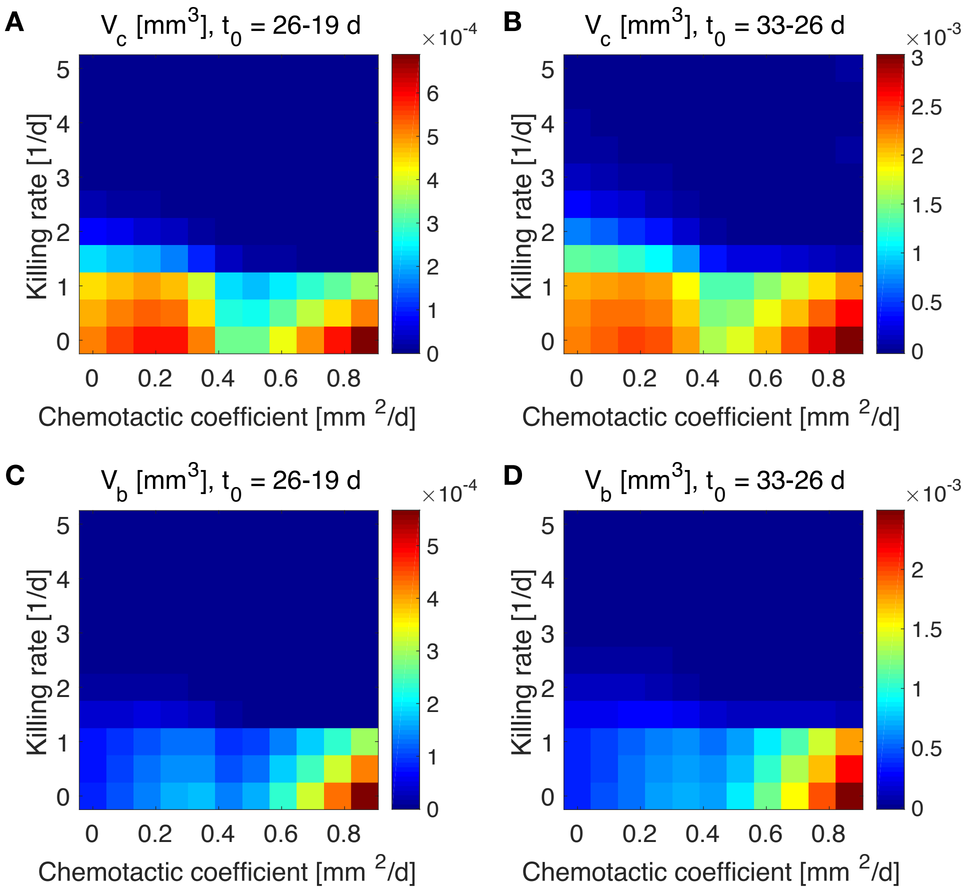


Figure S2: Influence of bacteria administration time $t_0$ on tumor and bacteria volumes at the end of the simulations (day 50). The maps show the difference between the volumes obtained using the parameters listed in the figure titles.

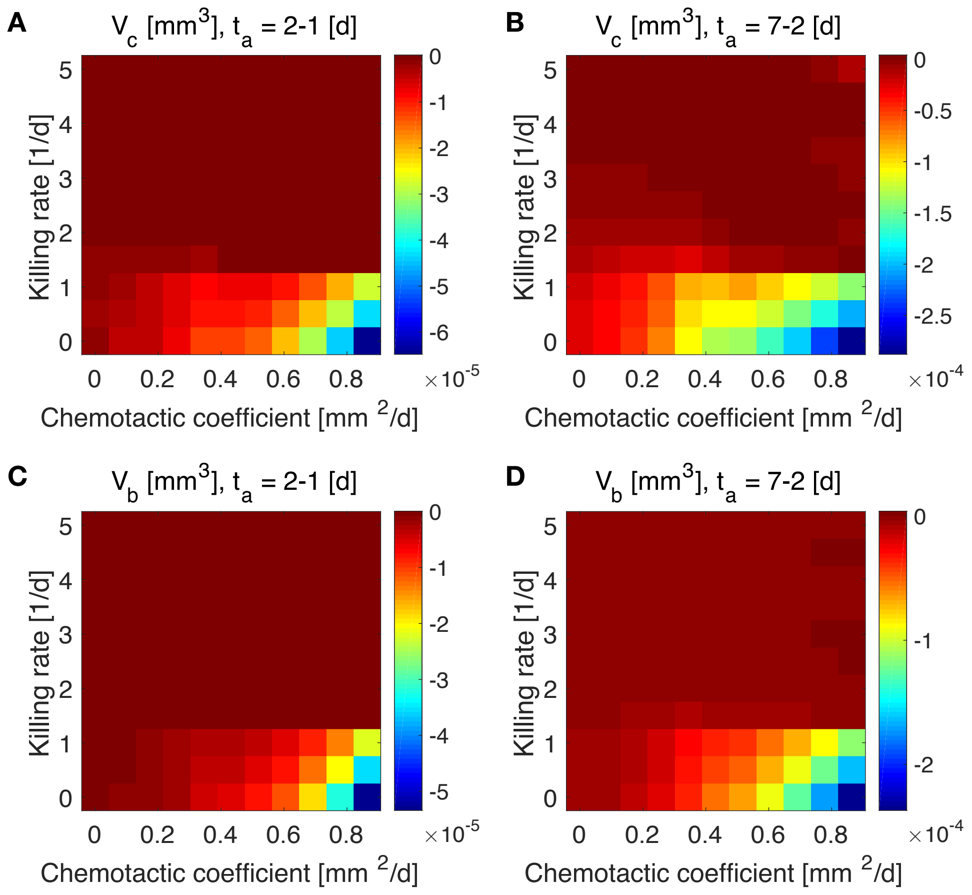


Figure S3: Influence of bacteria administration duration $t_{\mathrm{a}}$ on tumor and bacteria volumes at the end of the simulations (day 50). The maps show the difference between the volumes obtained using the parameters listed in the figure titles.

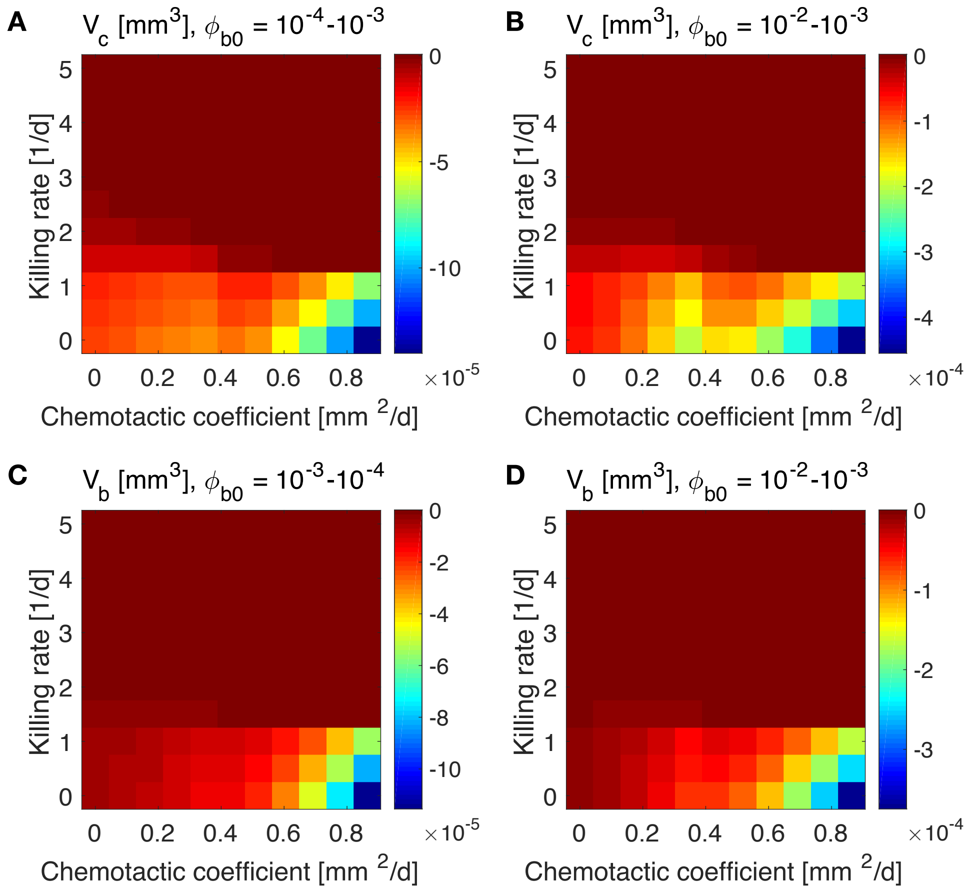


Figure S4: Influence of administered bacteria volume fraction $\phi_{b0}$ on tumor and bacteria volumes at the end of the simulations (day 50). The maps show the difference between the volumes obtained using the parameters listed in the figure titles.

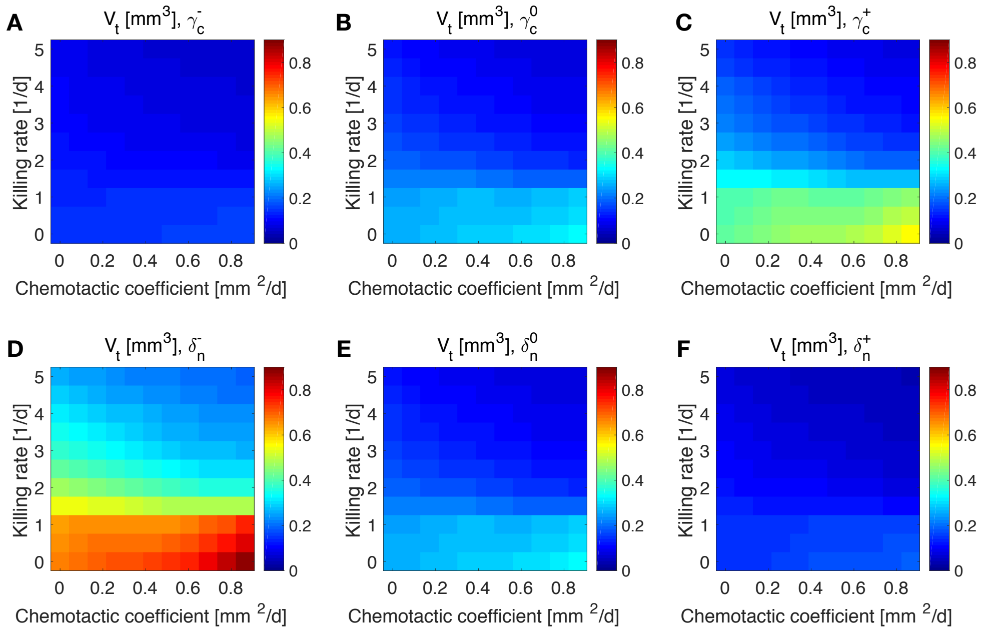


Figure S5: Final total spheroid volume for different parameters characterizing tumor cells. **A-C** Variation of tumor cell proliferation rate. **D-F** Variation of tumor cell oxygen consumption.
